# Multidimensional Associations between Cognition and Connectome Organization in Temporal Lobe Epilepsy

**DOI:** 10.1101/675884

**Authors:** Raúl Rodríguez-Cruces, Boris C. Bernhardt, Luis Concha

**Affiliations:** Universidad Nacional Autónoma de México, Instituto de Neurobiología Querétaro, Querétaro, México; MICA laboratory, Montreal Neurological Institute and Hospital, McGill University, Montreal Canada; Universidad Nacional Autónoma de México, Instituto de Neurobiología, Laboratory C-13, Boulervard Juriquilla 3001, Juriquilla, Querétaro, México. C.P. 76230, Querétaro, Querétaro, México

**Keywords:** Epilepsy, Connectome, Cognition, Multivariate, Network Neuroscience

## Abstract

**Objective:** Temporal lobe epilepsy (TLE) is known to affect large-scale structural networks and cognitive function in multiple domains. The study of complex relations between structural network organization and cognition requires comprehensive analytical methods and a shift towards multivariate techniques. The current work sought to identify multidimensional associations between cognitive performance and structural network topology in TLE.

**Methods:** We studied 34 drug-resistant TLE patients and 25 age- and sex-matched healthy controls. All participants underwent a comprehensive neurocognitive battery and multimodal MRI, allowing for large-scale connectomics, and morphological evaluation of subcortical and neocortical regions. Using canonical correlation analysis, we identified a multivariate mode that links cognitive performance to a brain structural network. Our approach was complemented by bootstrap-based clustering to derive cognitive subtypes and associated patterns of macroscale connectome anomalies.

**Results:** Both methodologies provided converging evidence for a close coupling between cognitive impairments across multiple domains and large-scale structural network compromise. Cognitive classes presented with an increasing gradient of abnormalities (increasing cortical and subcortical atrophy and less efficient white matter connectome organization in patients with increasing degrees of cognitive impairments). Notably, network topology characterized better the cognitive performance than morphometric measures. Thus, connectome characteristics featured as important markers of network reorganization and loss of inter-regional connectivity.

**Conclusions:** The multivariate approach emphasized the close interplay between cognitive impairment and large-scale network anomalies in TLE. Our findings contribute to understand the complexity of structural connectivity regulating the heterogeneous cognitive deficits found in epilepsy

## 1 Introduction

Temporal lobe epilepsy (TLE) is the most common drug-resistant epilepsy in adults and traditionally associated to mesiotemporal sclerosis, a lesion affecting the hippocampus and adjacent mesial structures[1]]. In addition to seizures, patients suffer from cognitive impairments that severely impact everyday functioning and wellbeing[2]]. In fact, TLE has traditionally been investigated by cognitive neuroscience as an important model to understand human memory and language dysfunction resulting from hippocampal damage[3].

Recent years have seen an evolution in our understanding of the cognitive landscape and structural compromise in TLE, fostered by an increasing administration of comprehensive neurocognitive phenotyping batteries and the advent of high-resolution and multimodal neuroimaging,[4, 5]. At the level of cognitive function, TLE is now recognized to perturb multiple domains not limited to memory and language processing[5, 6]. These findings are paralleled by mounting neuroimaging evidence suggesting diffuse grey and white matter abnormalities beyond the mesial temporal lobe, affecting a distributed network of cortical and subcortical structures as well as their connections[7–9].

While some studies have shown compromise of both white and grey matter regions in TLE patients relative to the degree of cognitive dysfunction[10–14], we lack a comprehensive understanding on the association between the extent of network reorganization and overall cognitive performance. Associations between brain structure and cognitive performance are likely complex, particularly when multiple metrics are used for neuroanatomical profiling on the one hand, and cognitive phenotyping on the other hand. Variable collinearities may furthermore challenge interpretability, and variables could lose their weight when tested individually. Multivariate analysis solves this problem by relating all measures in a single, compact mode[15]. Although converging evidence suggest an association between network organization and cognitive impairments in TLE[16], virtually no previous research leveraged multivariate techniques to identify salient brain cognition associations in the condition. It remains unknown if there is a structural white matter network pattern associated with the cognitive decline seen in patients. We hypothesize that whole brain structural network abnormalities seen in TLE are closely associated with the heterogeneous cognitive performance.

We examined the interplay between multidimensional cognitive performance and structural network compromise in TLE patients and healthy controls. All participants underwent state-of-the-art multimodal magnetic resonance imaging (MRI) and neurocognitive assays. Multivariate Canonical Correlation Analysis (CCA) evaluated associations between multi-domain cognitive impairment and whole brain structural connectome reorganization. These models were complemented by unsupervised clustering techniques to identify cognitive subtypes in the TLE cohort, for which we identified morphological and network-based signatures. We leveraged bootstrap-based stability assessments as well as cross-validation techniques to strengthen robustness and replicability of discovered network substrates of cognitive impairment. Finally, we made all code and data related to our study openly available.

## 2 Materials and Methods

### 2.1 Participants

The Ethics Committee of the Neurobiology Institute of the Universidad Nacional Autónoma de México approved this project (protocol code 019.H-RM) and written informed consent was obtained from all participants in the study according to the Declarations of Helsinki.

We recruited 34 adult ambulatory patients with drug-resistant TLE (Age= 29.7 *±* 11.1 years; 22 females) and 24 age- and sex-matched healthy controls (Age= 32.8 *±* 12.7 years; 18 females). Our cohort included 12 right TLE, 18 left TLE, and 4 bilateral TLE patients lateralized by seizure history and semiology, inter-ictal EEG recordings, and neuroimaging. All participants were right-handed native Spanish speakers. They did not have MRI contraindications nor other neurological comorbidities. Clinical features were obtained through a questionnaire-oriented interview upon referral (age at disease onset= 14.4 *±* 9.3 years; seizure frequency per month= 4.2 *±* 7.1, number of anti-epileptic drugs= 1.6 *±* 0.6, 35.2% had a history of febrile seizures).

### 2.2 Data acquisition

#### 2.2.1 Cognition

All participants underwent a comprehensive battery of cognitive tests: Wechsler Adult Intelligence Scale (WAIS-IV) and Wechsler Memory Scale (WMS-IV). We utilized the following index scores: auditory memory (AMI), visual memory (VMI), visual working memory (VWM), immediate memory (IMI), delayed memory (DMI), verbal comprehension (VCI), working memory (WMI), processing speed (PSI), and perceptual reasoning (PRI). All reported indices were normalized relative to a Mexican population and adjusted by age and education level. Details of the cognitive evaluation are described elsewhere[17].

#### 2.2.2 Magnetic Resonance Imaging

Images were acquired with a 3 Tesla Philips Achieva TX scanner with a 32-channel head coil. T1-weighted volumes (three-dimensional spoiled gradient echo) had a voxel resolution of 1×1×1 *mm*^3^, repetition time (TR) of 8.1 *ms*, echo time (TE) of 3.7 *ms*, flip angle of 8°, and field of view (FOV) dimensions of 179×256×256 *mm*^3^. Diffusion weighted images (DWI) were acquired with echo-planar imaging (EPI) and a 2×2×2 *mm*^3^ voxel resolution, TR=11.86 *s* and TE=64.3 *ms*, and FOV=256×256×100 *mm*^3^. DWI were sensitized to 60 different diffusion gradient directions (b=2000 *s/mm*^2^); one b=0 *s/mm*^2^ volume was also acquired. An additional b=0 *s/mm*^2^ volume was obtained with reversed phase encoding polarity to account for geometric distortion corrections.

### 2.3 Image processing

#### 2.3.1 Diffusion MRI processing

a. *Diffusion weighted volumes* (DWI) were denoised via data redundancy criteria from linear dimensinality reduction, followed by non-uniform intensity normalization. Reverse phase encoding from two b=0 *s/mm*^2^ volumes was used to estimate and correct for geometric distortions. DWI volumes were linearly registered to the b=0 *s/mm*^2^ images for motion correction and diffusion gradient vectors were rotated according to the transformation matrix.
b. *Structural connectome parameterization*. Using FreeSurfer v5.3.0, MRtrix 3.0, and FSL 5.0.6, individual structural connectivity matrices were calculated based on the corrected DWI and using Spherical-deconvolution Informed Filtering of Tractograms, SIFT[18] with anatomically constrained tractography models ACT[19]. A total of 162 nodes were defined merging the cortical parcellation from the Destrieux Atlas and volBrain’s subcortical segmentation (**Supplementary Table 1**). Whole brain tractography was first calculated using ACT with 20 million streamlines seeded on the gray-white matter interface, with maximum deviation angle of 22.5°, maximum length of 250 mm, minimum length of 10 mm. Tractograms were filtered via SIFT to 2 million streamlines (Fig. 1 **top left**). Connection weights between nodes (*N*_*SIFT*_) was defined as the streamline counts following SIFT[20, 21], (Fig. 1 **top right**). This procedure has shown high reproducibility in structural connectomics ([22]). **Figure 1:**
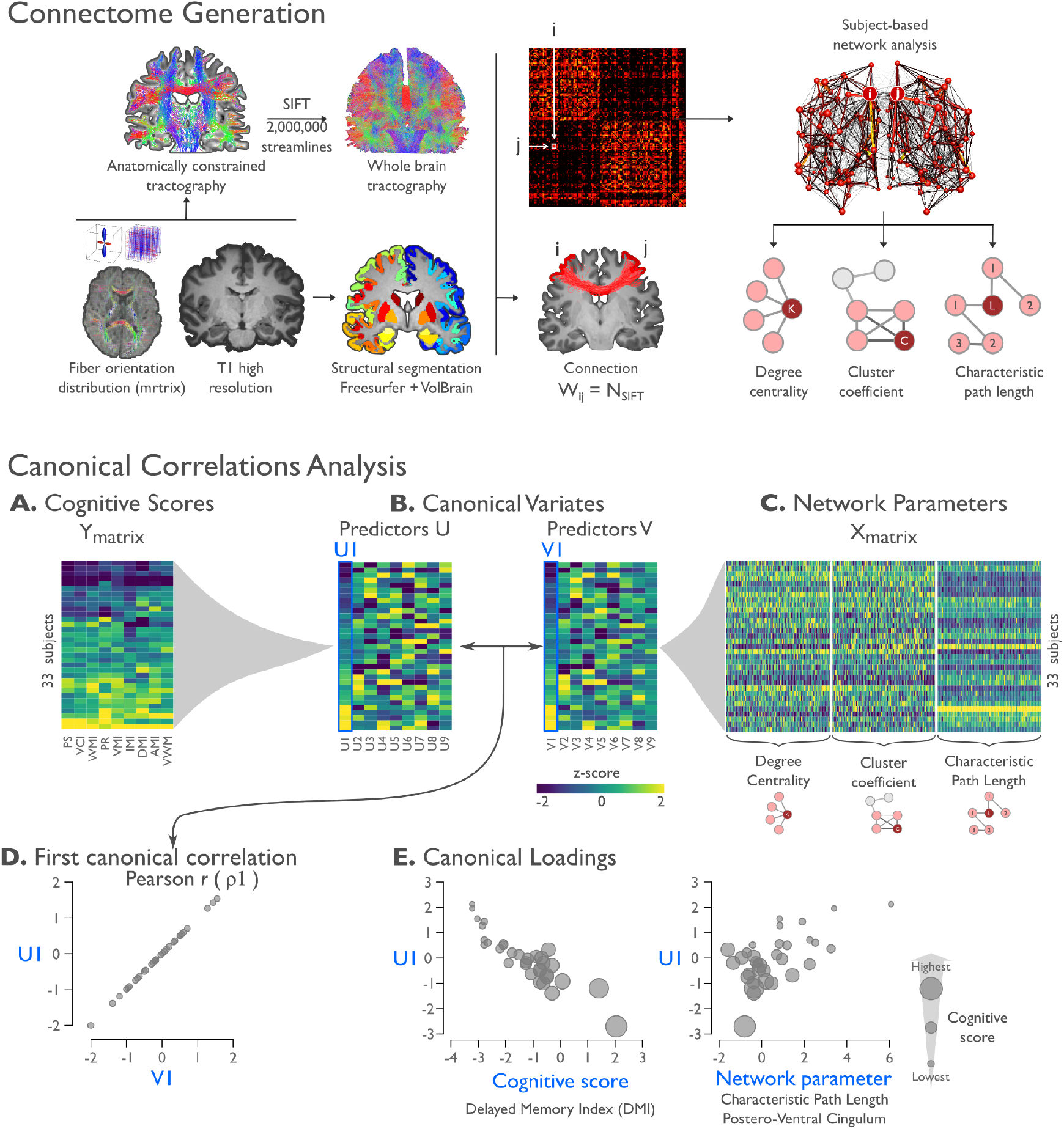
Connectome generation. **Top left:** Whole-brain connectomes were built using mrtrix, based on streamline counts derived from anatomically constrained tractography and spherical deconvolution informed filtering of tractograms (SIFT). Nodes were defined by merging the cortical segmentation of Destrieux Atlas and Volbrain’s subcortical segmentation. Connection weight W_ij_ was defined as the streamline count between two nodes ij following SIFT. **Top right:** To study network topology, degree centrality, clustering coefficient, and characteristic path length were calculated based on the adjacency matrices. Cluster coefficient was calculated using the Onnela algorithm. **Multivariate analysis: canonical correlations analysis. A.** For each participant, the cognitive scores, excluding IQ were combined into matrix Y. Similarly, the nodal network measurements associated with a brain region were concatenated to a matrix X (**panel C**). **B.** The canonical variates are synthetic predictors (V and U) that maximize the correlations between the cognitive scores and the network parameters. **D.** The correlation between the first canonical variate U1 and V1 is referred as the first canonical correlation ρ1. **E.** The canonical loadings measure the linear correlation between an original variable of the cognitive scores Y_j_ or the network parameters X_j_ and a canonical variate.

Connectivity matrices were analyzed using the *igraph R package* (igraph.org/r). We focused on path length, clustering coefficient, and degree centrality, the most widely used graph-theoretical parameters in the TLE literature ([23]), also given that these measurements offer a compact description of global network topology and local connectivity embedding ([24]). We computed the *clustering-coefficient (C)* as a measure of segregation, which provides information about the level of local connections in a network. Additionally, we measured the characteristic *path length (L)* as a measure of network integration with short path lengths indicating globally efficient networks. Dijkstra algorithm was used to calculate the inverse distance matrix and infinite path lengths were replaced with the maximum finite length. Finally, we calculated *degree centrality (k)* to characterize the relevance of the individual nodes. We set the appropriate threshold for network metrics with recursive thresholds from 0.1-0.9 until reaching convergence stability (Supplementary Fig.1). We set the appropriate threshold for network metrics with recursive thresholds from 0.1-0.9 until reaching convergence stability (**Supplementary Fig. 1**).

#### 2.3.2 Structural MRI processing

a. *Hippocampal volumetry*. The T1-weighted volumes were processed using volBrain (volbrain.upv.es), which provides automated patch-based hippocampal and subcortical delineation with high accuracy in controls and TLE patients. Hippocampi were individually inspected by a trained rater, and hippocampal volumes were normalized by intracranial volume.
b. *Cortical thickness analysis*. Cortical thickness was measured for each participant using FreeSurfer v5.3.0. T1 images were pre-processed through non-local-means denoising and N4 bias field correction prior to FreeSurfer segmentation. After processing, pial and white matter surfaces were visually inspected by a qualified trained rater and corrected if necessary. Individual surfaces were registered to a surface template with 20,484 surface points (fsaverage5) and a surface-based Gaussian diffusion filter with a full width at half maximum of 20mm was applied, similar to our previous studies[25].

### 2.4 Multivariate Analysis

#### 2.4.1 Regularized canonical correlation analysis

Canonical correlation analysis (CCA) assessed multivariate associations between cognitive scores and structural connectome measures (Fig. 1 **bottom**). Unlike principal components analysis (PCA) that reduces the number of variables in one set to components that emphasize variation in the data, CCA investigates the overall correlation between two multivariate datasets. CCA was recently employed in a large cohort of healthy adults to identify associations between neuroimaging-based connectivity measures on the one hand, and lifestyle, demographic, and psychometric measures on the other hand[26].

First, we built a CCA to evaluate associations between connectome-derived parameters (k, C, and L) of all brain regions, and cognitive performance. Network parameters were concatenated into one row vector per subject, resulting in a matrix X (subjects x *network parameters*). We excluded IQ because of its high correlation with all the remaining scores, resulting in a matrix Y (subjects x *cognitive scores*). The main objective of CCA is to estimate canonical variates (U and V) that maximize the correlation between *network parameters*-X and *cognitive scores*-Y (Fig. 1B **bottom** and **Supplementary Fig. 2A**). Resulting canonical variates can be ordered (*U*_1_ − *U*_*n*_, and *V*_1_ − *V*_*n*_), with the first explaining the largest proportion of covariance among sets X-Y.

Additionally, canonical loadings represent the relationship between an original variable and a canonical variate (Fig. 1E **bottom**). As the number of subjects was less than the number of variables in both data sets, we included two regularization parameters, estimated via leave-one-out cross-validation with recursive search (**Supplementary Fig. 2B**). These parameters were employed to reduce overfitting due to the large number of variables[27]. Each CCA model was permuted and bootstrapped 10,000 times to estimate confidence intervals and significance.

In addition to the main TLE-CCA model, we evaluated the following additional models to test for specificity: one with morphological measures (i.e., volumetric of subcortical and cortical areas), one including only controls, one controlling matrix X and Y for hippocampal volume and mean cortical thickness, and a full model that included network parameters, clinical features and volumes. The latter was performed in order to reveal the possible clinical contributions additional to the structural parameters in the definition of cognitive profile.

#### 2.4.2 Stable cluster analysis for cognitive phenotypes

Clustering techniques have been suggested to capture heterogeneity in different clinical cohorts, and applied to cognitive variables in epileptic groups([4, 5]). We performed a classification based in the cognitive scores of TLE patients to identify associations between our multivariate analysis and cognitive performance. Robust cognitive phenotypes were identified via unsupervised and bootstrap-supported analysis to identify maximally stable clusters (Fig. 2, [28]).

**Figure 2:**
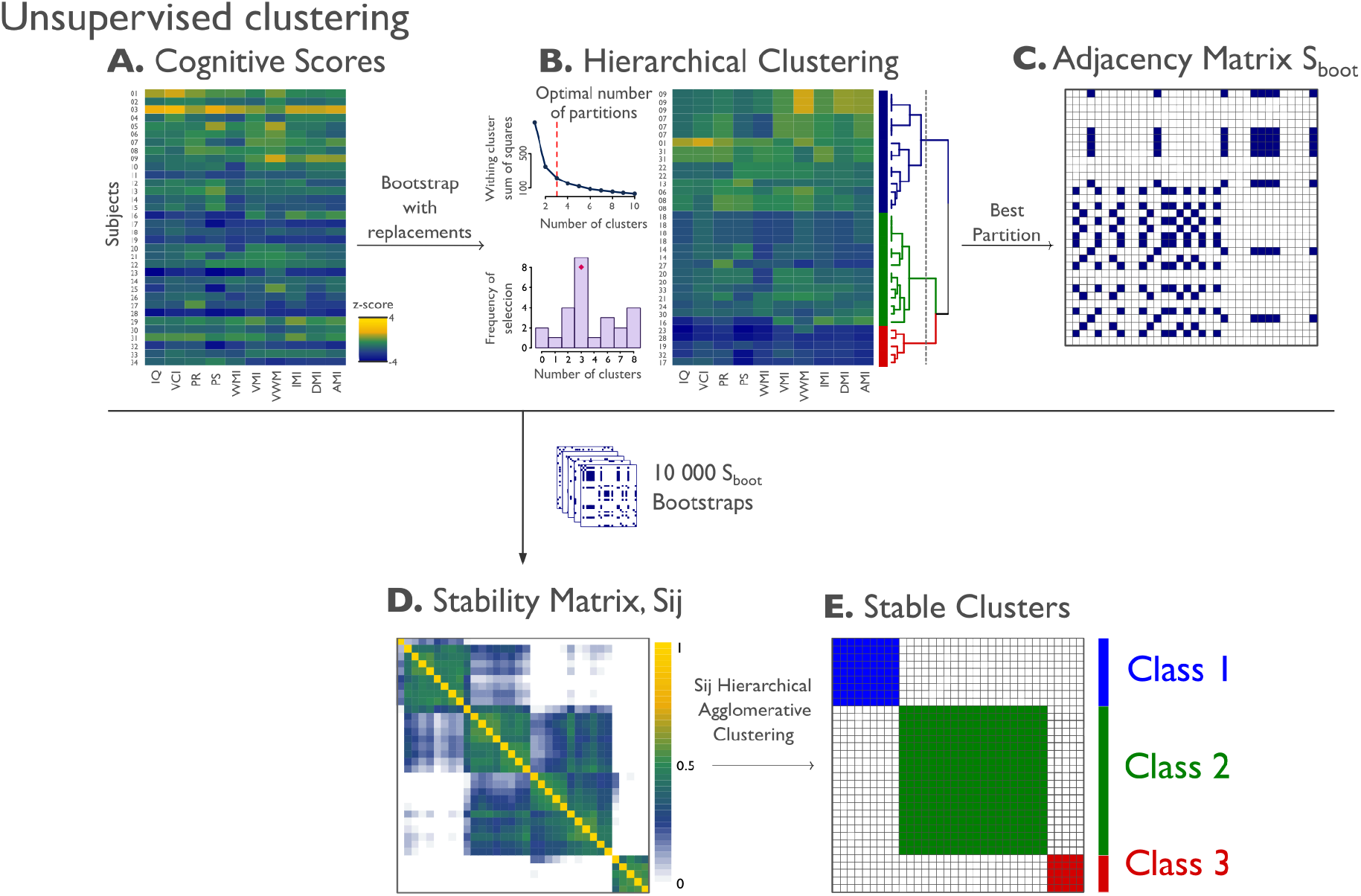
Unsupervised clustering. **A)** Cognitive features as z-scores with respect to controls are shown for each patient (rows). **B)** Example of a bootstrap with replacements realization with Ward D2 hierarchical agglomerative clustering. The optimal number of clusters (k) was determined from k=2-30 [29]. **C)** Adjacency matrix of the optimal partition for each bootstrap S_boot_, where S_boot_ij equals 1 if participants i and j belong to the same partition and 0 otherwise. **D)** After 10000 bootstraps, final stability matrix S_ij_ that represents the percentage of times a subject i was classified similarly to subject j. **E)** Hierarchical agglomerative clustering is performed over the stability matrix S_ij_ matrix, clustering converges on a three-subtype solution in our cohort.

#### 2.4.3 Class difference analysis

Feature data including hippocampal and subcortical volumes as well as cortical thickness, were z-scored relative to controls and sorted into ipsilateral/contralateral with respect to the seizure focus ([30]).

1. *Clinical variables* were compared between classes using ANOVAs followed by Tukey’s post-hoc correction for multiple comparisons.
2. *Topological complex network*. Each nodal parameter (k, C, L) was sorted into ipsilateral/contralateral relative to hemispheric TLE lateralization and compared to controls for each Class and represented as effect size (Cohen’s D). For statistical comparison a node-level (ROI) t-test was performed for each TLE class compared to controls. Differences in nodal network parameters were corrected for multiple comparisons at a two-tailed false discovery rate (FDR) of q=0.025.
3. *Cortical thickness and subcortical volumes* were compared to controls, and corrected with the mean cortical thickness for each subject. Surface-based analysis was done using SurfStat[31]. Effect size of the cortical thickness (Cohen’s D) between group differences was calculated for each Class, and compared to controls at a vertex level using t-tests, and corrected for multiple comparisons with FDR, q<0.025.

## 3 Results

### 3.1 Multivariate association analyses

Canonical correlation analysis revealed one significant association between cognitive and structural connectome features in TLE (permutation-test p<0.05; Fig. 3). Associated patterns of loadings showed that reduced cognitive scores were related to reduced degree centrality and clustering, together with increased path length. Network loadings encompassed measures from both cortical and subcortical regions and were high in both ipsilateral and contralateral regions. Specifically, longer path lengths were related to lower cognitive scores in TLE, indicating associations between lower global efficiency and worse cognitive performance. Similarly, reduced degree centrality was found in bilateral superior frontal lobes, and precentral gyrus. Finally, clustering coefficient in ipsilateral parietal and middle frontal gyrus related to lower cognitive scores. When clinical and volume features were added to the CCA, results were consistent with the original model, however negative loadings related years of study and volume of both hippocampi with lower cognitive scores (**Supplementary Fig. 3**).

**Figure 3:**
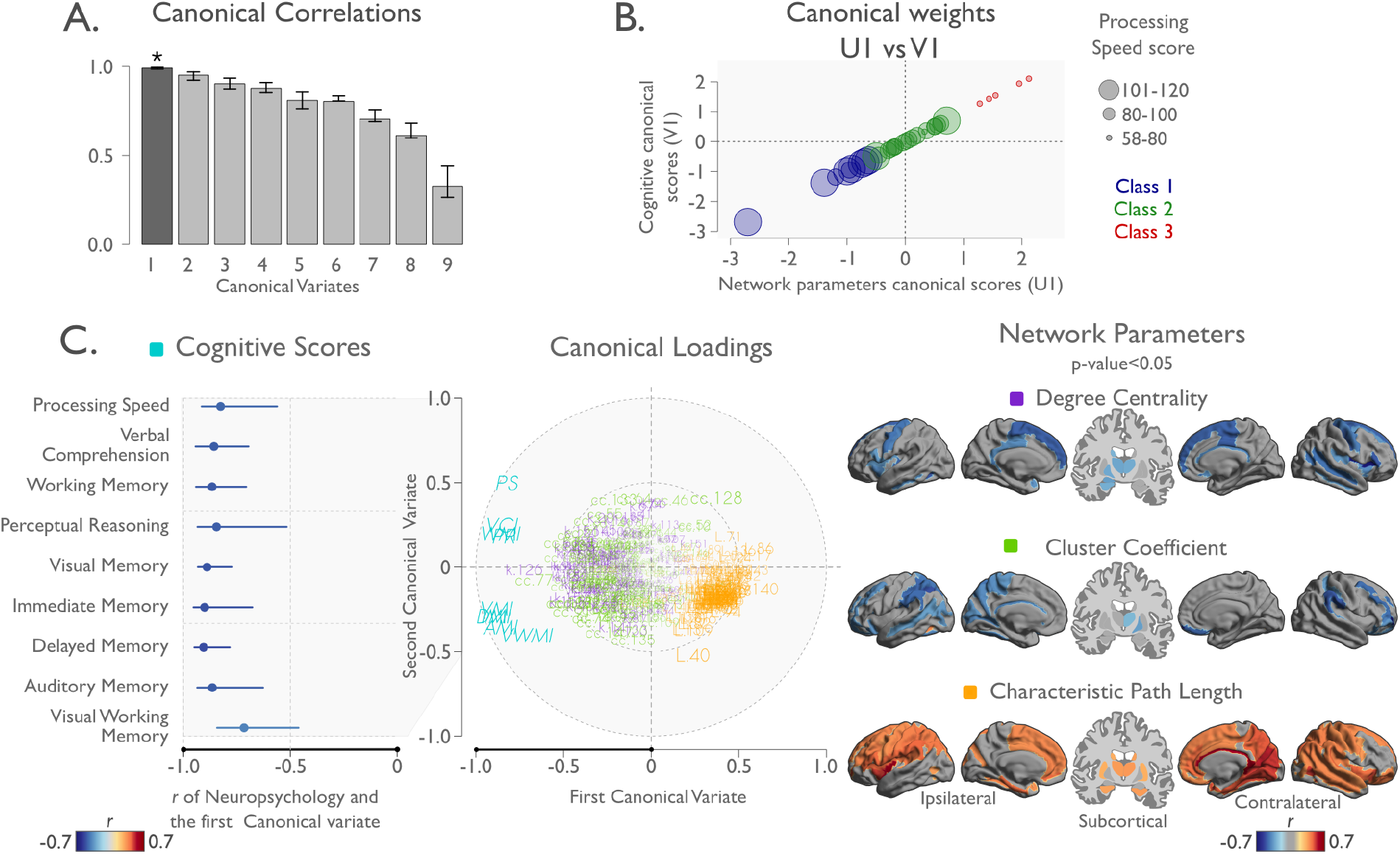
Regularized canonical correlation solution. **A.** Canonical correlations for each canonical variate, each with confidence interval and significance (**\*** and darker gray indicate statistical significance). **B.** Scatterplot of the canonical weights assigned to the cognitive scores against the network parameter of the first canonical variate for each TLE patient (U1 *versus V*1). Processing speed score (PS) is shown as size of the circles, and color represents cognitive Class. **C.** Canonical cross-loadings of the first and second canonical variates for the cognitive scores and network parameters. Loadings were obtained by correlating each of the variables directly with a canonical variate. **C-Left** panel shows the correlation between each cognitive score and the first canonical variate. The lines represent the confidence interval over the first canonical variate (x-axis). **C-Middle** panel shows the cognitive scores and network loadings on the plane of the first and second canonical variates. Network loadings are represented by color: Purple for degree, green for cluster coefficient and orange for characteristic path length. Cognitive loadings are represented in cyan: AMI-Auditory memory, VMI-visual memory, VWM-visual working memory, IMI-immediate memory, DMI-delayed memory, VCI-verbal comprehension, WMI-working memory, PS-processing speed and PR-perceptual reasoning. **C-Right** panel shows the significant network loadings of the first canonical variate, projected to the surface space and split by network measurement.

Multivariate CCA between morphological measures and cognitive characteristics did not yield any significant associations in patients (**Supplementary Fig. 4**). Likewise, none of the canonical correlations were significant in controls (**Supplementary Fig. 5**). Furthermore, the topological measures were independently associated with cognitive performance when controlling for hippocampal atrophy and cortical thickness (**Supplementary Fig. 6**)

### 3.2 Cognitive Classes

Bootstrap-based clustering of cognitive profiles converged on three cognitive classes in our TLE cohort (Fig. 2.E, Fig. 4.A). Cognitive deficits showed an increasing gradient over the three Classes, yet the pattern of these deficits was specific for each. Patients in Class 1 had cognitive scores within normal range, those in Class 2 showed mild impairment in memory-specific domains, and patients in Class 3 displayed pronounced impairment across all domains, with prominent reduction of processing speed (Table 1).

**Table 1:** Clinical data by Class

|  | Class 1 | Class 2 | Class 3 |
| --- | --- | --- | --- |
| *Number* | 9 | 20 | 5 |
| *HS % presence* | **0.33\*** | **0.45\*** | **0.80\*** |
| *Gender % female* | 0.56 | 0.75 | 0.40 |
| *Age years* | 28.7 (10.8) | 30.9 (12.2) | 26.4 (7) |
| *Education years* | 14.3 (2.9) | 12.3 (2.7) | **8.4 (1.3)\*<sup>1,2</sup>** |
| *Age at onset years* | 19.2 (12.2) | 13.8 (7.3) | 8 (7.1) |
| *Duration years* | 9.4 (8.7) | 17.1 (14.7) | 18.4 (8.3) |
| *AED* | 1.3 (0.5) | 1.7 (0.7) | 1.6 (0.6) |
| ***Global network parameters*** |  |  |  |
| *Degree centrality* | 93.6 (3.6) | 89.3 (5.6) | 87.1 (4.3) |
| *Path length $\times 10^{-4}$* | 31.2 (2.8) | 31.3 (3.2) | **38.6 (11.6)\*<sup>1,2</sup>** |
| *Cluster coefficient* | 0.72 (0.01) | 0.71 (0.01) | 0.71 (0.01) |
| ***Cognitive performance*** |  |  |  |
| *Intelligence quotient* | 101.0 (13.7) | **82.8 (7.6)\*<sup>1</sup>** | **63.6 (14.7)\*<sup>1,2</sup>** |
| *Verbal comprehension* | 100.7 (20.6) | **84.4 (7.8)\*<sup>1</sup>** | **64.6 (13.1)\*<sup>1,2</sup>** |
| *Working memory* | 100.2 (11.6) | **82.3 (9.4)\*<sup>1</sup>** | **63.6 (8.8)\*<sup>1,2</sup>** |
| *Perceptual reasoning* | 103.9 (8.9) | **86.5 (8.7)\*<sup>1</sup>** | **65.2 (7.8)\*<sup>1,2</sup>** |
| *Processing speed* | 101.9 (10.4) | **90.4 (10.3)\*<sup>1</sup>** | **66.2 (9.6)\*<sup>1,2</sup>** |
| *Auditory memory* | 97.1 (18.4) | **78.2 (11.0)\*<sup>1</sup>** | **49.2 (2.6)\*<sup>1,2</sup>** |
| *Visual working memory* | 101.9 (7.7) | **76.3 (13.5)\*<sup>1</sup>** | **52.0 (7.3)\*<sup>1,2</sup>** |
| *Immediate memory* | 99.1 (15.1) | **75.5 (11.6)\*<sup>1</sup>** | **44.6 (4.6)\*<sup>1,2</sup>** |
| *Delayed memory* | 96.4 (16.6) | **73.7 (12.7)\*<sup>1</sup>** | **49.2 (3.3)\*<sup>1,2</sup>** |

AED: number of antiepileptic drugs, HS, hippocampal sclerosis. Age, education, onset, duration, AEDs, degree centrality, path length, and cluster coefficient shown as mean (standard deviation). \* Significant difference compared to controls ( $p_{adjusted} < 0.05$ ). Superscripts indicate significant difference with respect to the Class indicated by the number ( $p_{adjusted} < 0.05$ ).

**Figure 4:**
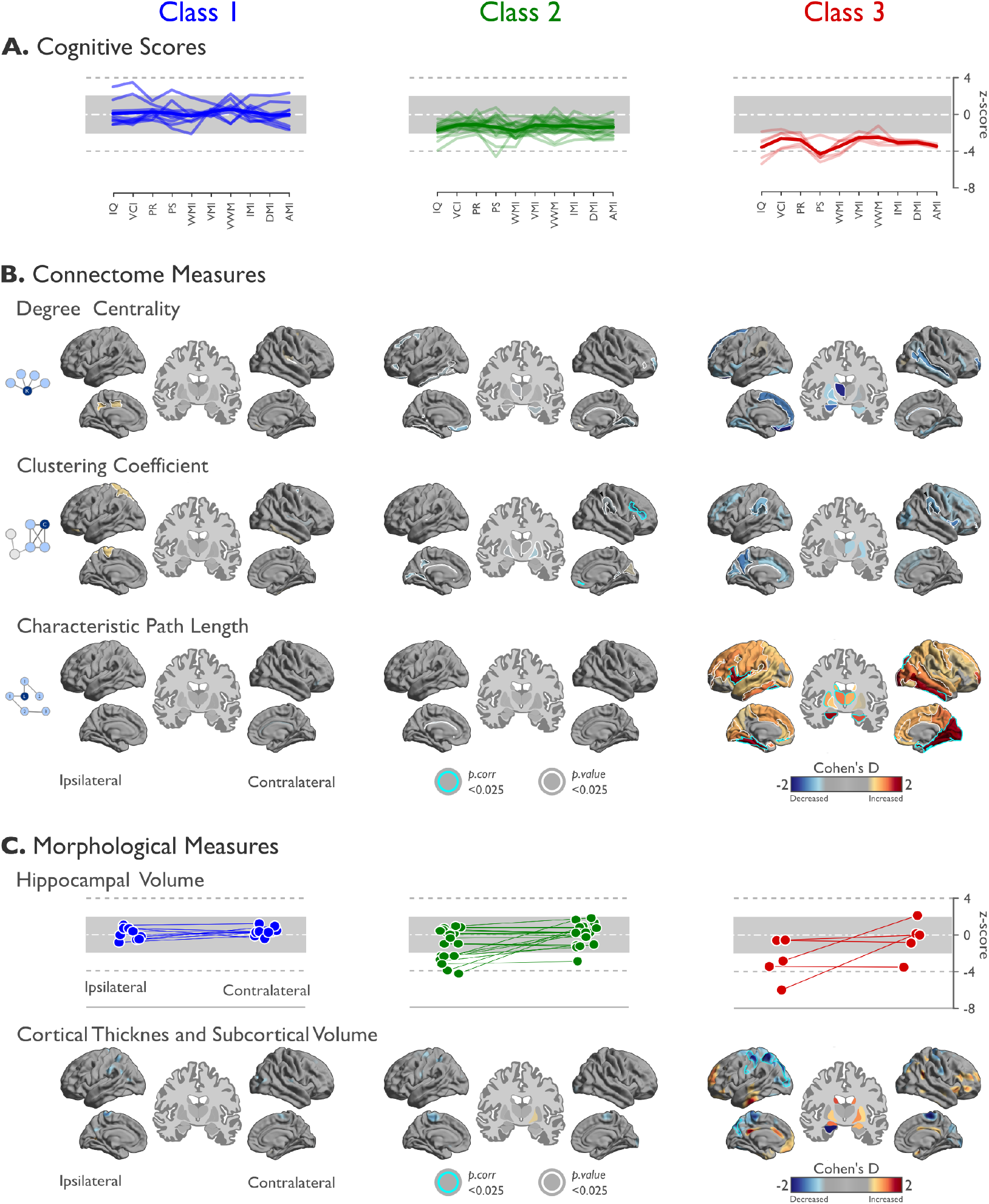
Differences by cognitive class. **A.** Cognitive scores for TLE patients by cognitive Class. Each patient is represented as a line indicating their normalized cognitive scores based on control, and the mean of each Class represented as a thick line. **B.** Connectome measures. For each metric, effect size (Cohen’s D) of each Class compared to controls is projected on cortical surfaces. Significant differences corrected for multiple comparisons are outlined in cyan; white outlines represent uncorrected p<0.025. **C.** Morphological Measures. Hippocampal volumes were z-scored relative to controls. Group differences in cortical thickness and subcortical volume shown as Cohen’s D effect sizes. Thickness is relative to the mean vertex value of each Class, while volume is the mean volume of each subcortical region.

Class 1 patients presented with older age of epilepsy onset, more years of education, and the shortest disease duration. Despite of these clinical differences, overall findings were similar when controlling for age, duration of epilepsy, and number of antiepileptic drugs. Hippocampal sclerosis was less prevalent in Class 1 (33%) than Class 3 (80%, Table 1).

### 3.3 Connectome-level and morphological compromise across cognitive Classes

Gradual network organization abnormalities were observed across Classes with most marked changes in Class 3, intermediate differences in Class 2, and only subtle changes in Class 1 (Fig. 4). Class 1 presented restrained increases of degree centrality and clustering coefficient relative to controls in (ipsilateral) cingulate and parietal cortices at uncorrected thresholds (Fig. 4.B). Class 2 showed significantly decreased clustering in the contralateral suborbital sulcus and inferior frontal sulcus (*p*_*FDR*_ < 0.025). At a connectome-wide level, Class 3 showed the most marked increases of characteristic path length while Classes 1 and 2 were rather normal (*p*_*FDR*_ < 0.025). In Class 3, path length increases were most marked in the lateral and medial temporal lobes in both hemispheres, the ipsilateral frontal and the contralateral occipital lobe.

Similar to the findings in network parameters, structural MRI markers showed an increasing gradient of changes from Class 1 (the most similar to controls) to Class 3 (the most abnormal, Fig. 4.C). Hippocampal volumes in Class 1 were within the control range, while Class 2 and 3 had and increasing degrees of hippocampal atrophy. Cortical thinning was also most pronounced in Class 3, particularly in parietal areas ipsilateral to the focus.

A final integrative qualitative analysis examined associations between the rCCA and clustering solutions. This analysis revealed a tight relation between the first canonical variate (U1) with our robust clustering solution for all cognitive scores (Fig. 5). When we controlled our CCA model for hippocampal volume ipsilateral to the lesion and mean cortical thickness, the main canonical loadings were preserved, but the canonical weights lost their hierarchical relation with the cognitive metrics.

**Figure 5:**
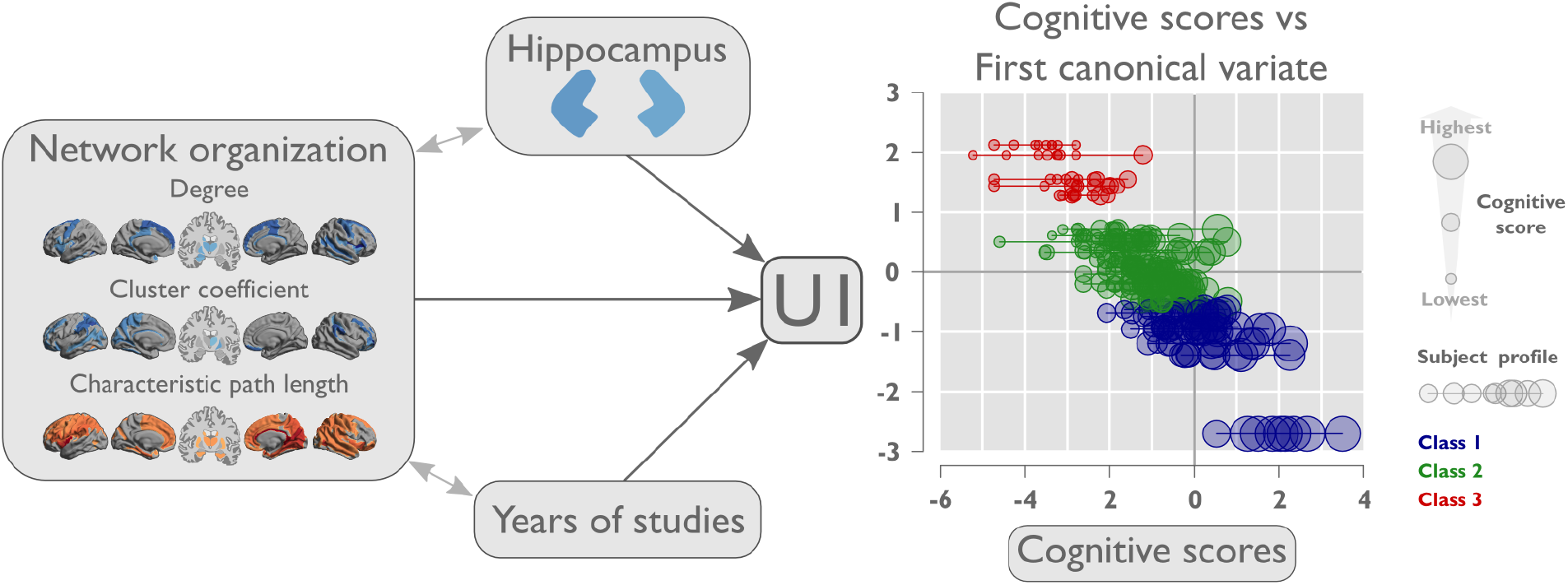
Cognitive convergence between the first canonical variate and cognitive scores. Plot of the relation between the first canonical variate (U1) and all the cognitive scores, colored by class. Y-axis represents the value of the first canonical variate of the rCCA-TLE model for each subject, while on x axis we plot all the cognitive scores as z-score based on controls. Each subject’s cognitive profile is shown as a horizontal line. The size of circles represents the score for each cognitive test. Individual cognitive tests are not distinguishable in this plot.

## 4 Discussion

The current work targeted the complex interplay between structural connectome reorganization and cognition in patients with drug-resistant temporal lobe epilepsy (TLE). Harnessing two complementary multivariate data science methodologies (*i.e.*, canonical correlation analysis and data-driven clustering) we observed converging evidence for a close link between the overall degree of white matter network perturbations and multi-domain cognitive impairment in our patients. In particular, we found less efficient network organizations in patients with more marked cognitive difficulties. Notably, although complementary cortical thickness analysis revealed marked morphological anomalies in the same patient cohort, these measures were less closely associated to cognitive dysfunction than white matter connectome metrics. Furthermore, associations were less marked in healthy controls, suggesting disease specificity. Overall, these findings provide novel and robust evidence for a close and specific coupling of cognitive phenotypes and white matter connectome topology in patients with temporal lobe epilepsy, suggesting a network level pattern underlying broad variations in cognitive function seen in these patients.

Core to our data acquisition was a multidomain cognitive phenotyping together with a whole-brain neuroimaging and connectomics paradigm. The use of a broad neuropsychological battery instead of restricted psychometric testing was motivated by prior observations suggesting that TLE impacts not only language and memory, but rather a diverse set of cognitive domains also including attentional and executive functioning[4, 5]. Similarly, we employed hippocampal volumetry, cortical thickness analyses, as well as diffusion MRI connectomics to assess macroscale brain anomalies in both grey and white matter compartments. Prior histopathological and morphological studies have indeed suggested that although TLE is generally associated to mesiotemporal anomalies[1], it is rarely associated to a confined focal pathological substrate[32, 33]. Instead, an increasing number of MRI-based cortical thickness assessments and subcortical shape analyses have indicated a rather distributed structural compromise, often characterized by bilateral temporo-limbic as well as fronto-central atrophy ([7, 9, 25]). Similarly, a growing body of white matter tractographic analyses and network neuroscience work leveraging graph theoretical formalisms of structural connectomes suggested atypical white matter organization and microstructure not limited to the temporal lobe, but in a rather widespread topographic distribution radiating outwards from the mesiotemporal epicenter[10, 14, 34, 35]. Although these distributed abnormalities have been hypothesized to affect cognitive function[2], there are so far only sporadic systematic attempts to relate imaging measures to multidomain cognitive phenotypes in TLE. In fact, among those studies associating structural anomalies and cognitive performance in TLE[11–14, 17, 36], the majority has been rather selective, focusing on the relation between specific brain measures on the one hand, and specific cognitive domains on the other hand.

We harnessed multivariate associative techniques as well as bootstrap-based clustering to integrate the broad panorama of cognitive phenotypes in TLE with a comprehensive array of structural neuroimaging measures. The former class of models[15], in our case a regularized canonical correlation analysis (CCA), provides a set of sparse components capturing complex covariation patterns between network parameters and cognitive profiles. A recent study leveraged CCA to identify gradual associations between functional connectome configurations and a wide array of factors related to lifestyle, demographics and psychometric function in a large cohort of young healthy adults, describing a positive-negative mode of co-variation between observable behavior and self-report measures and functional connectome organization[26]. In our TLE cohort, CCA revealed a consistent pattern of associations characterized by distributed increases in connectome path length related to reduced cognitive performance. Previous reports have shown similar increases of characteristic path length in this condition compared to controls, suggesting overall reduced global network efficiency[8, 16, 37, 38]. Further elements of the brain-behavior covariation mode encompassed low frontal lobe clustering coefficient together with reduced parietal hubness in patients with reduced cognitive functions, potentially indicating a breakdown of frontal and parietal network segregation that may ultimately reflect network level consequences secondary to microstructural anomalies previously reported in these systems, potentially indicative of axonal damage, myelin alteration, as well as reactive astrogliosis[39]. As multivariate associative techniques like CCA can overfit, we incorporated several additional elements to ensure specificity and robustness. Firstly, we verified consistency via cross-validation techniques, dispelling potential hyper-optimization of within-sample associations at the expense to out-of-sample generalization. Secondly, we did not observe similar associations in controls, suggesting specificity to TLE. Finally, associations were more marked at the level of white matter connectomes than for grey matter morphometry, confirming overall a close association between white matter connectome architecture and cognitive phenotypes in the condition.

Further support for the consistency of the brain-behavior association in our patients was provided by data-driven clustering of the cognitive profiles, additionally supported in the current work using bootstrap based stability maximization[28]. Subtyping of epileptic patients based on cognitive profiles has previously been employed to identify a spectrum of cognitive function[4, 5, 17, 36]. The applied method converged on a three-class solution with gradual cognitive impairments and overall corresponding degrees of brain anomalies, assuring that cognitive impairment in TLE is indeed related to an increased load of white matter connectome reorganization, together with hippocampal and cortical grey matter atrophy. Integrative analyses confirmed that these discovered cognitive classes provide a different viewpoint on the dimensional multivariate mode of covariation seen via CCA (Fig. 5). Of note, the prevalence of hippocampal atrophy increased across the three cognitive classes, with the class showing the most marked cognitive dysfunction and connectome anomalies (*i.e.*, Class 3) also presenting the highest degree of hippocampal volume loss. Conversely, TLE laterality was similarly distributed across classes, potentially due to the broader range of domains evaluated in the current study than in work focusing on language and/or memory, which generally support more marked impairment in left compared to right TLE[40].

In addition to the novel use of advanced multivariate techniques and state-of-the-art connectomics and cognitive phenotyping, our findings are well anchored in overarching assumptions on the link between brain structure and function in healthy and diseased brains. Our findings encourage the use of multivariate methods and contribute to understand the complexity of structural connectivity regulating the heterogeneous cognitive deficits found in epilepsy.

## 5 Acknowledgements

We sincerely thank the patients and their families, as well as our control subjects, for their willingness to participate, the medical specialists who helped us with their recruitment, and the clinical personnel at the National Laboratory for magnetic resonance imaging. We are grateful to Juan Ortíz-Retana, Erick Pasaye and Leopoldo González-Santos for technical assistance. We extend our gratitude to the many people who have at some point participated in this study, performing patient recruitment, scanning, and clinical evaluations: Leticia Velázquez-Pérez, David Trejo, Héctor Barragán, Arturo Domínguez, Ildefonso Rodríguez-Leyva, Ana Luisa Velasco, Luis Octavio Jiménez, Daniel Atilano, Elizabeth González Olvera, Rafael Moreno, Vicente Camacho, Ana Elena Rosas, and Alfonso Fajardo.

## 8 Data avaliability statement

Phenotypic and imaging data, as well as code for statistical analysis are freely available on our OSF repository: https:/doi.org/10.17605/OSF.IO/JBDN2. For processing details, please see: https://github.com/rcruces/cognition_conectomics_TLE. For additional information and further details about all rCCA models, connectome parameterization, and the long table of ROIs of our segmentations, please see our OSF repository and supplementary material.

SurfStat is available via http://mica-mni.github.io/surfstat.

