## Supplementary File for "Multidimensional Associations between Cognition and Connectome Organization in Temporal Lobe Epilepsy"

Github Code: https://github.com/rcruces/cognition_conectomics_TLE

OSF Data: DOI 10.17605/OSF.IO/JBDN2

#### Contents

**List of Figures** ii

**List of Tables** ii

**1 Methods: Density thresholds of network metrics** 1

**2 Methods: Stable cluster analysis for cognitive phenotypes** 2

**3 Methods: Regularized canonical correlation analysis** 3

**4 Results: Regularized canonical correlation analysis** 4

**5 Table: Cortical and subcortical segmentation** 8

**References** 12

### 1 Methods: Density thresholds of network metrics

To assess the stability of network parameters, a proportional threshold was calculated for each subject connectivity matrix by preserving the proportion of the strongest weights. All the other weights and the main diagonal were set to 0. Network parameters (k, C, L) were recursively thresholded from 0.1-0.9 to find the appropriate threshold for network metrics reaching convergence stability.

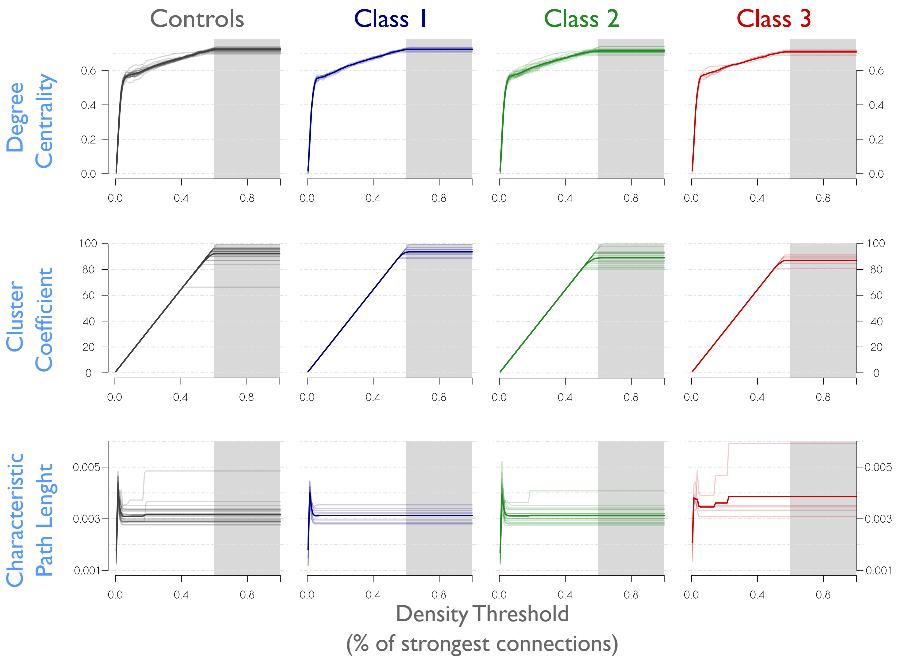

Supplementary Figure 1: The plots represents the different network metrics as a function of network density (percentage of strongest connection). Columns are the groups and row the featured parameters. Each subject is shown as a line and the mean of each class is shown as a thicker line. Density thresholds converge on a stable parameter over 60% of the connections for all features. Characteristic path length stabilizes at 30%.

#### 2 Stable cluster analysis for cognitive phenotypes

Unsupervised clusterization was based on Bellec et al. 1. In brief, we took the dataset of cognitive scores and TLE lateralization as a numeric input matrix Y (**Fig. 2A**). Bootstrap with replacements was then applied to Y and we obtained a new set of samples of the matrix Y, with each realization defined as Yboot. Next, we applied Ward D2 hierarchical agglomerative clustering to each Yboot (**Fig. 2B**), and the optimal number of clusters (k) was determined from k=1-30 [2]. The next step consisted in the generation of the best partition adjacency matrix (Sboot), where (Sbootij) = 1 if participants i and j belong to the same partition and 0 otherwise (**Fig. 2C**). After 10,000 bootstraps, the clustering stability was quantified as a matrix Sij, where each pair of subjects (ij) represents the probability of belonging to the same group (**Fig. 2D**). To obtain stable clusters, hierarchical agglomerative clustering (HAC) was applied to Sij using the k with highest probability from the distribution of all partitions obtained from all bootstraps. Bootstrap based clustering converged on a 3-subtype solution in our TLE cohort (**Fig. 2E**).

### 3 Regularized canonical correlation analysis

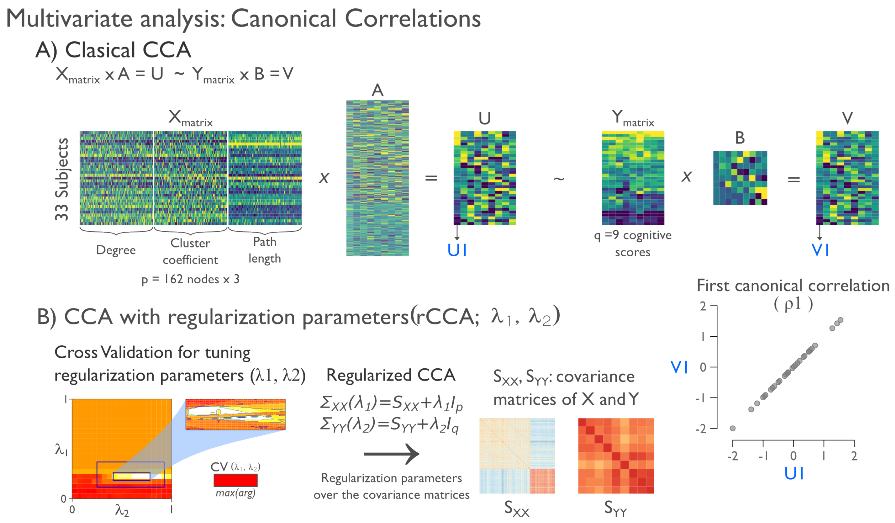

Supplementary Figure 2: **Multivariate analysis: Canonical correlations. A.** Diagram of the regularized canonical correlations analysis. For each participant, the nodal network measurements associated with a brain region were concatenated to a single row vector of matrix $X$. Similarly, cognitive scores were combined into matrix $Y$. These two sets of variables were used for the rCCA, that estimates the matrices $A$ and $B$, which maximize the correlation between the linear relation of each column vector of $V$ and $U$. The resulting variables $U_1$ and $V_1$ are the first canonical variate and their correlation is referred as the first canonical correlation $\rho_1$. Columns in $A$ describe the extent to which that canonical variate is related to a network parameter of a specific brain region, and columns in $B$ describe the extent to which that canonical variate is present in a particular cognitive score. The canonical loadings measure the linear correlation between an original variable of $X_j$ or $Y_j$ and the selected canonical variate. **B.** The regularization parameters ($\lambda 1$, $\lambda 2$ for $X$ and $Y$) are applied to the covariance matrices ($S_{XX}$, $S_{YY}$). A cross-validation procedure is performed to determine optimal regularization parameters. Cross-validation with recursive search over a grid from 0 to 1 for regularization parameters tuning optimization. The search region enclosed in the blue square is seen enlarged on the right.

### 4 Regularized canonical correlation analysis

We performed several rCCA models in order to prove that our results from the main rCCA in TLE patients were due to the disease. We explored the rCCA in controls, TLE controlling both sets X and Y for hippocampal volume and mean cortical thickness, a full model of TLE including clinical variables and hippocampal volume, and finally a rCCA model in TLE using only the ROI volumes controlled for intracraneal volume.

All figures have the same distribution of three panels A to C:

**A.** Canonical correlations with confidence interval and significance (* and lighter gray). **B.** Scatterplot of the canonical weights assigned to the neuropsychological against the network parameter of the first canonical variate for each subject ($U1$ versus $V1$). The size of the point represents the rank of the Processing speed score (PS) and the color is the cognitive class belonging. **C.** Canonical cross-loadings of the first and second canonical variate for the neuropsychometrical scores and the nodal network parameters. The loadings are obtained by correlating each of the variables directly with a canonical variate. **C.Left panel** shows the correlation between each neuropsychological score and the first and second canonical variate. The lines represent the confidence interval over the first canonical variate (x-axis). The font size shows the variance explained of each variable and the color is associated with the correlation value of the neuropsychological canonical loading with the first canonical variate. **C.middle** represents the neuropsychological and nodal network loadings on the plane of the first and second canonical variate. The nodal network loadings are represented by color. Purple for the degree, green for the cluster coefficient and orange for the characteristic path length. Neuropsychological loadings are represented in cyan: AMI-Auditory memory, VMI-visual memory, VWM-visual working memory, IMI-immediate memory, DMI-delayed memory, VCI-verbal comprehension, WMI-working memory, PS-processing speed and PR-perceptual reasoning. **C. Right panel** shows only the significant nodal network loadings of the first canonical variate, projected to the surface space and split by network measurement. The strength of the correlations is represented with the color gradient from gray to purple, green or orange, for degree, cluster coefficient or characteristic path length, respectively.

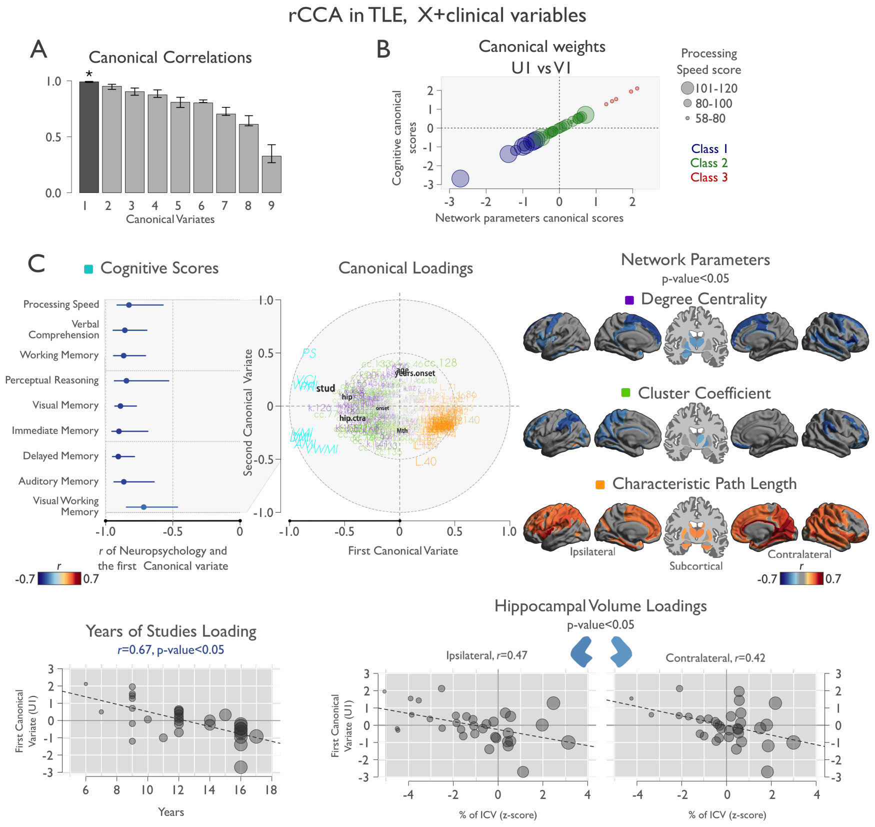

Supplementary Figure 3: rCCA in TLE, X + clinical variables + mean cortical thickness + hippocampal volume. rCCA model of TLE with clinical variables did not show any difference in the loadings of the network parameters when compared to the original rCCA model of TLE. Middle panel C shows the localization of the clinical loadings with respect to the first and second canonical variate (black font, stud: years of studies, hip: ipsilateral hippocampus, hip.ctra: contralateral hippocampus, onset: age at onset, Mth: mean cortical thickness, Age: current age, years.onset: years since onset). Panel C bottom shows the relation between the first canonical variate and the significant variables; years of studies and both hippocampal volumes. Circle size represents the value of the variable in the X-axis

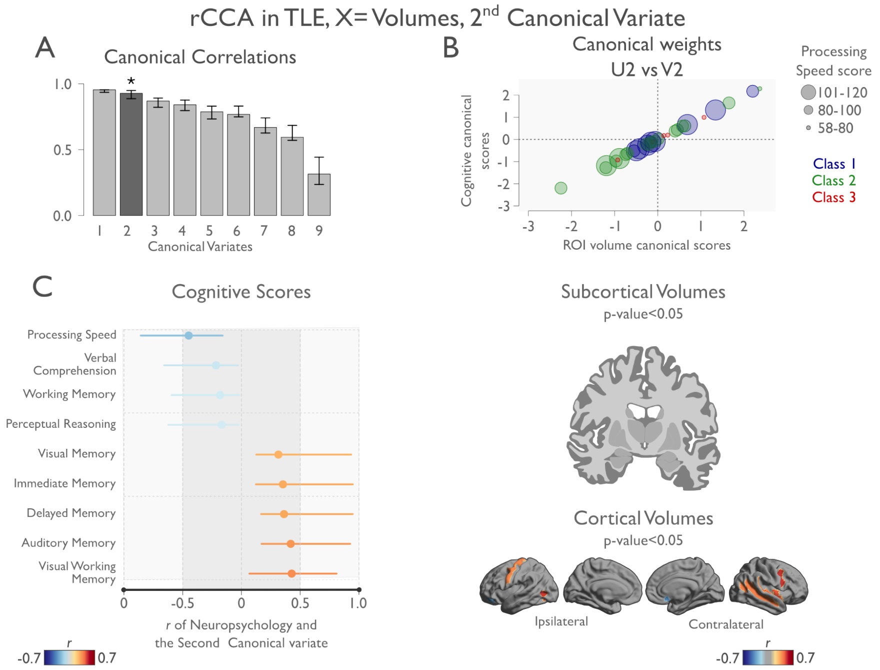

Supplementary Figure 4: rCCA model of TLE patients, were the X set consists in the volumes of all cortical and subcortical regions of interest controlled by intracaraneal volume. Only the second canonical variate was significant, but shows no strong relationships with neuropsychology as in the original rCCA TLE model of network parameters.

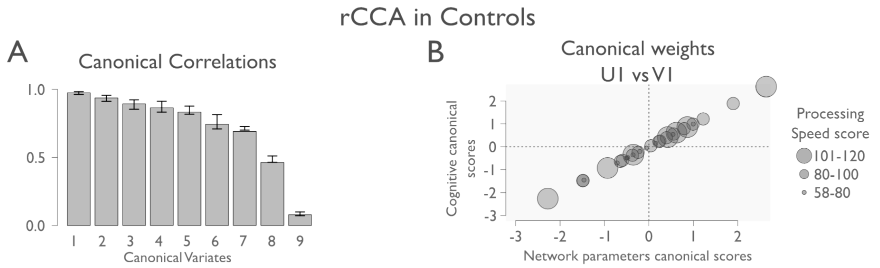

Supplementary Figure 5: The rCCA model of controls did not show any significant canonical correlation.

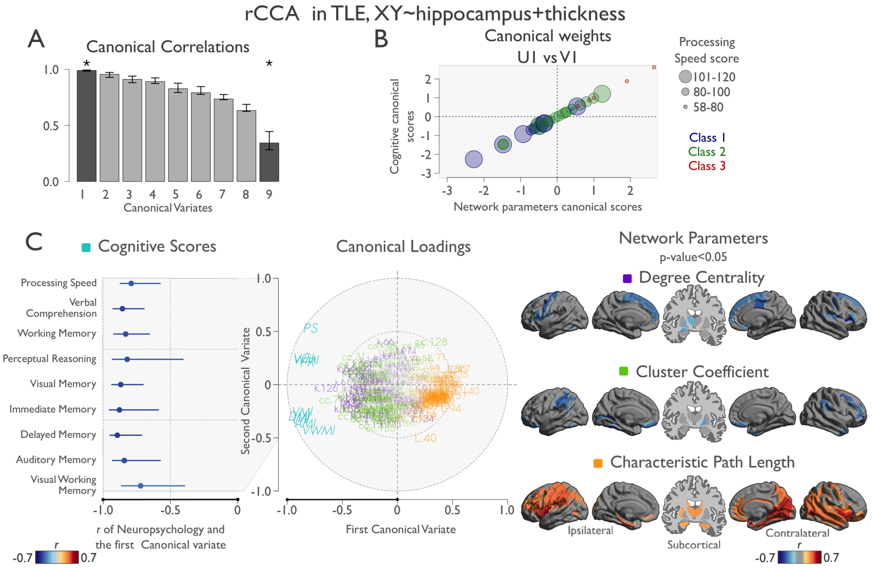

Supplementary Figure 6: rCCA model of TLE patients, where the data sets of neuropsychological and structural variables (x and Y) are controlled by the volume of hippocampus ipsilateral to the seizure onset and the mean cortical thickness value.

### 5 Cortical and subcortical segmentation

Nodes were defined by merging the cortical segmentation of Destrieux Atlas and Volbrain's subcortical segmentation [3, 4].

Table 1: Subject based segmentation

| ROI ID | ROI name | Side | Full name |
|---|---|---|---|
| 1 | L.F.mar | left | G_and_S_frontomargin |
| 2 | L.O.inf | left | G_and_S_occipital_inf |
| 3 | L.par.C | left | G_and_S_paracentral |
| 4 | L.sub.C | left | G_and_S_subcentral |
| 5 | L.tran.F.pol | left | G_and_S_transv_frontopol |
| 6 | L.Cin.A | left | G_and_S_cingul-Ant |
| 7 | L.Cin.MA | left | G_and_S_cingul-Mid-Ant |
| 8 | L.Cin.MP | left | G_and_S_cingul-Mid-Post |
| 9 | L.Cin.PD | left | G_cingul-Post-dorsal |
| 10 | L.Cin.PV | left | G_cingul-Post-ventral |
| 11 | L.CU | left | G_cuneus |
| 12 | L.F.inf.Op | left | G_front_inf-Opercular |
| 13 | L.F.inf.Or | left | G_front_inf-Orbital |
| 14 | L.F.inf.Tr | left | G_front_inf-Triangul |
| 15 | L.F.mid | left | G_front_middle |
| 16 | L.F.sup | left | G_front_sup |
| 17 | L.Ins.ig | left | G_Ins_lg_and_S_cent_ins |
| 18 | L.Ins.sh | left | G_insular_short |
| 19 | L.O.mid | left | G_occipital_middle |
| 20 | L.O.sup | left | G_occipital_sup |
| 21 | L.OT.latfus | left | G_oc-temp_lat-fusifor |
| 22 | L.OT.medling | left | G_oc-temp_med-Lingual |
| 23 | L.OT.medparhip | left | G_oc-temp_med-Parahip |
| 24 | L.Orb | left | G_orbital |
| 25 | L.P.infang | left | G_pariet_inf-Angular |
| 26 | L.P.infsupmar | left | G_pariet_inf-Supramar |
| 27 | L.P.sup | left | G_parietal_sup |
| 28 | L.pos.C | left | G_postcentral |
| 29 | L.pre.C | left | G_precentral |
| 30 | L.pre.CU | left | G_precuneus |
| 31 | L.R | left | G_rectus |
| 32 | L.sub.Call | left | G_subcallosal |
| 33 | L.T.sup.Gttra | left | G_temp_sup-G_T_transv |
| 34 | L.T.sup.lat | left | G_temp_sup-Lateral |
| 35 | L.T.sup.pol | left | G_temp_sup-Plan_polar |
| 36 | L.T.sup.tem | left | G_temp_sup-Plan_tempo |
| 37 | L.T.inf | left | G_temporal_inf |
| 38 | L.T.mid | left | G_temporal_middle |

Table 1: Subject based segmentation

| ROI ID | ROI name | Side | Full name |
|---|---|---|---|
| 39 | L.Lat.FisHor | left | Lat_Fis-ant-Horizont |
| 40 | L.Lat.FisVer | left | Lat_Fis-ant-Vertical |
| 41 | L.Lat.FisPos | left | Lat_Fis-post |
| 43 | L.pole.O | left | Pole_occipital |
| 44 | L.poleT | left | Pole_temporal |
| 45 | L.S.Cal | left | S_calcarine |
| 46 | L.S.Cen | left | S_central |
| 47 | L.S.cingM | left | S_cingul-Marginalis |
| 48 | L.S.Ins.ant | left | S_circular_insula_ant |
| 49 | L.S.Ins.inf | left | S_circular_insula_inf |
| 50 | L.S.Ins.sup | left | S_circular_insula_sup |
| 51 | L.S.coll.tra.ant | left | S_collat_transv_ant |
| 52 | L.S.coll.tra.pos | left | S_collat_transv_post |
| 53 | L.S.F.inf | left | S_front_inf |
| 54 | L.S.F.mid | left | S_front_middle |
| 55 | L.S.F.sup | left | S_front_sup |
| 56 | L.S.IPJ | left | S_interm_prim-Jensen |
| 57 | L.S.Iptra | left | S_intrapariet_and_P_trans |
| 58 | L.S.O.mid.Lu | left | S_oc_middle_and_Lunatus |
| 59 | L.S.O.sup.tra | left | S_oc_sup_and_transversal |
| 60 | L.S.O.ant | left | S_occipital_ant |
| 61 | L.S.O.lat | left | S_oc-temp_lat |
| 62 | L.S.OT.med.ling | left | S_oc-temp_med_and_Lingual |
| 63 | L.S.Orb.lat | left | S_orbital_lateral |
| 64 | L.S.Orb.med | left | S_orbital_med-olfact |
| 65 | L.S.Orb.H | left | S_orbital-H_Shaped |
| 66 | L.S.PO | left | S_parieto_occipital |
| 67 | L.S.per.Call | left | S_pericallosal |
| 68 | L.S.pos.C | left | S_postcentral |
| 69 | L.S.pre.C.inf | left | S_precentral-inf-part |
| 70 | L.S.pre.C.sup | left | S_precentral-sup-part |
| 71 | L.S.sub.Orb | left | S_suborbital |
| 72 | L.S.sub.P | left | S_subparietal |
| 73 | L.S.T.inf | left | S_temporal_inf |
| 74 | L.S.T.sup | left | S_temporal_sup |
| 75 | L.S.T.tra | left | S_temporal_transverse |
| 77 | R.F.mar | right | G_and_S_frontomargin |
| 78 | R.O.inf | right | G_and_S_occipital_inf |
| 79 | R.par.C | right | G_and_S_paracentral |
| 80 | R.sub.C | right | G_and_S_subcentral |
| 81 | R.tran.F.pol | right | G_and_S_transv_frontopol |
| 82 | R.Cin.A | right | G_and_S_cingul-Ant |
| 83 | R.Cin.MA | right | G_and_S_cingul-Mid-Ant |

Table 1: Subject based segmentation

| ROI ID | ROI name | Side | Full name |
|---|---|---|---|
| 84 | R.Cin.MP | right | G_and_S_cingul-Mid-Post |
| 85 | R.Cin.PD | right | G_cingul-Post-dorsal |
| 86 | R.Cin.PV | right | G_cingul-Post-ventral |
| 87 | R.CU | right | G_cuneus |
| 88 | R.F.inf.Op | right | G_front_inf-Opercular |
| 89 | R.F.inf.Or | right | G_front_inf-Orbital |
| 90 | R.F.inf.Tr | right | G_front_inf-Triangul |
| 91 | R.F.mid | right | G_front_middle |
| 92 | R.F.sup | right | G_front_sup |
| 93 | R.Ins.ig | right | G_Ins_lg_and_S_cent_ins |
| 94 | R.Ins.sh | right | G_insular_short |
| 95 | R.O.mid | right | G_occipital_middle |
| 96 | R.O.sup | right | G_occipital_sup |
| 97 | R.OT.latfus | right | G_oc-temp_lat-fusifor |
| 98 | R.OT.medling | right | G_oc-temp_med-Lingual |
| 99 | R.OT.medparhip | right | G_oc-temp_med-Parahip |
| 100 | R.Orb | right | G_orbital |
| 101 | R.P.infang | right | G_pariet_inf-Angular |
| 102 | R.P.infsupmar | right | G_pariet_inf-Supramar |
| 103 | R.P.sup | right | G_parietal_sup |
| 104 | R.pos.C | right | G_postcentral |
| 105 | R.pre.C | right | G_precentral |
| 106 | R.pre.CU | right | G_precuneus |
| 107 | R.R | right | G_rectus |
| 108 | R.sub.Call | right | G_subcallosal |
| 109 | R.T.sup.Gttra | right | G_temp_sup-G_T_transv |
| 110 | R.T.sup.lat | right | G_temp_sup-Lateral |
| 111 | R.T.sup.pol | right | G_temp_sup-Plan_polar |
| 112 | R.T.sup.tem | right | G_temp_sup-Plan_tempo |
| 113 | R.T.inf | right | G_temporal_inf |
| 114 | R.T.mid | right | G_temporal_middle |
| 115 | R.Lat.FisHor | right | Lat_Fis-ant-Horizont |
| 116 | R.Lat.FisVer | right | Lat_Fis-ant-Vertical |
| 117 | R.Lat.FisPos | right | Lat_Fis-post |
| 119 | R.pole.O | right | Pole_occipital |
| 120 | R.poleT | right | Pole_temporal |
| 121 | R.S.Cal | right | S_calcarine |
| 122 | R.S.Cen | right | S_central |
| 123 | R.S.cingM | right | S_cingul-Marginalis |
| 124 | R.S.Ins.ant | right | S_circular_insula_ant |
| 125 | R.S.Ins.inf | right | S_circular_insula_inf |
| 126 | R.S.Ins.sup | right | S_circular_insula_sup |
| 127 | R.S.coll.tra.ant | right | S_collat_transv_ant |

Table 1: Subject based segmentation

| ROI ID | ROI name | Side | Full name |
|---|---|---|---|
| 128 | R.S.coll.tra.pos | right | S_collat_transv_post |
| 129 | R.S.F.inf | right | S_front_inf |
| 130 | R.S.F.mid | right | S_front_middle |
| 131 | R.S.F.sup | right | S_front_sup |
| 132 | R.S.IPJ | right | S_interm_prim-Jensen |
| 133 | R.S.Iptra | right | S_intrapariet_and_P_trans |
| 134 | R.S.O.mid.Lu | right | S_oc_middle_and_Lunatus |
| 135 | R.S.O.sup.tra | right | S_oc_sup_and_transversal |
| 136 | R.S.O.ant | right | S_occipital_ant |
| 137 | R.S.O.lat | right | S_oc-temp_lat |
| 138 | R.S.OT.med.ling | right | S_oc-temp_med_and_Lingual |
| 139 | R.S.Orb.lat | right | S_orbital_lateral |
| 140 | R.S.Orb.med | right | S_orbital_med-olfact |
| 141 | R.S.Orb.H | right | S_orbital-H_Shaped |
| 142 | R.S.PO | right | S_parieto_occipital |
| 143 | R.S.per.Call | right | S_pericallosal |
| 144 | R.S.pos.C | right | S_postcentral |
| 145 | R.S.pre.C.inf | right | S_precentral-inf-part |
| 146 | R.S.pre.C.sup | right | S_precentral-sup-part |
| 147 | R.S.sub.Orb | right | S_suborbital |
| 148 | R.S.sub.P | right | S_subparietal |
| 149 | R.S.T.inf | right | S_temporal_inf |
| 150 | R.S.T.sup | right | S_temporal_sup |
| 151 | R.S.T.tra | right | S_temporal_transverse |
| 152 | L.thal | Left | Thalamus-Proper |
| 153 | L.caud | Left | Caudate |
| 154 | L.putm | Left | Putamen |
| 155 | L.pall | Left | Pallidum |
| 156 | L.hipp | Left | Hippocampus |
| 157 | L.amyg | Left | Amygdala |
| 158 | L.accm | Left | Accumbens-area |
| 159 | R.thal | Right | Thalamus-Proper |
| 160 | R.caud | Right | Caudate |
| 161 | R.putm | Right | Putamen |
| 162 | R.pall | Right | Pallidum |
| 163 | R.hipp | Right | Hippocampus |
| 164 | R.amyg | Right | Amygdala |
| 165 | R.accm | Right | Accumbens-area |
